## Supplementary Material for "Crosslinking by ZapD drives the assembly of short FtsZ filaments into toroidal structures in solution"

1- Department of Cellular and Molecular Biophysics, Max Planck Institute of Biochemistry, 82152 Martinsried, Germany.
2- Department of Molecular Structural Biology, Max Planck Institute of Biochemistry, 82152 Martinsried, Germany.
3- Exzellenzcluster ORIGINS, Boltzmannstr. 2, 85748 Garching, Germany.
4- Molecular Interactions Facility, Centro de Investigaciones Biológicas Margarita Salas, Consejo Superior de Investigaciones Científicas (CSIC), 28040 Madrid, Spain.
5- Cryo-EM facility, Max Planck Institute of Biochemistry, 82152 Martinsried, Germany.
6- Centro de Investigaciones Biológicas Margarita Salas, Consejo Superior de Investigaciones Científicas (CSIC), 28040 Madrid, Spain.
7- Helmholtz Pioneer Campus, Helmholtz Munich, 85764 Neuherberg, Germany; Department of Chemistry, Technical University of Munich, 85748 Garching, Germany.

This file contains

- Legends to Supplementary Movies 1-3
- Supplementary Figures S1-S16
- Supplementary Table 1

### Legends to Supplementary Movies

***Supplementary Movie 1.*** Tomogram of a FtsZ toroidal structure promoted by ZapD showed in Figure 3a followed by segmentation of the tomogram. FtsZ filaments are in blue. Successive rotations of the segmented volume allow to visualize the structure of the toroid in 3D.

***Supplementary Movie 2.*** Isosurface from the toroid showed in Figure 2a. FtsZ filaments are shown in blue and putative ZapD connections in magenta. A close-up view and rotations of the segmented volume show the filament meshwork and the connections by ZapD in 3D.

***Supplementary Movie 3.*** Tomogram of a FtsZ straight bundle formed at high concentration of ZapD proteins showed in Figure 5b. Successive segmentation of the tomogram with FtsZ filaments labelled in blue and putative ZapD connections in magenta. Rotations, close-up views help the interpretation of the data and show the 3D structure of the straight bundle.

a

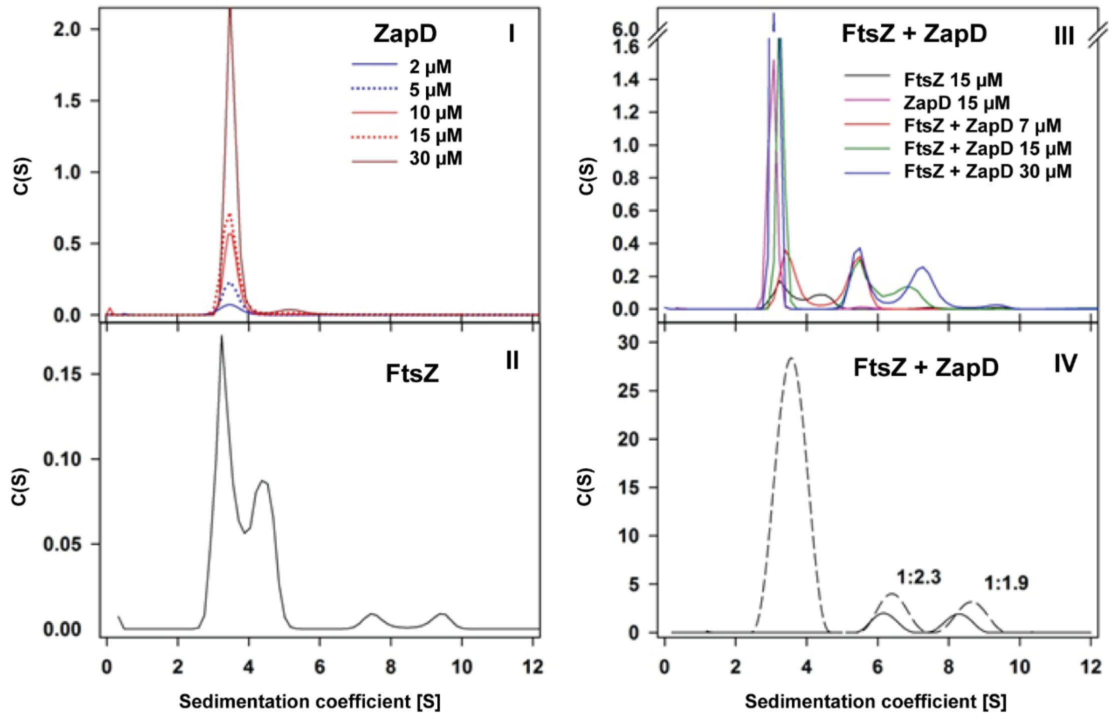


b

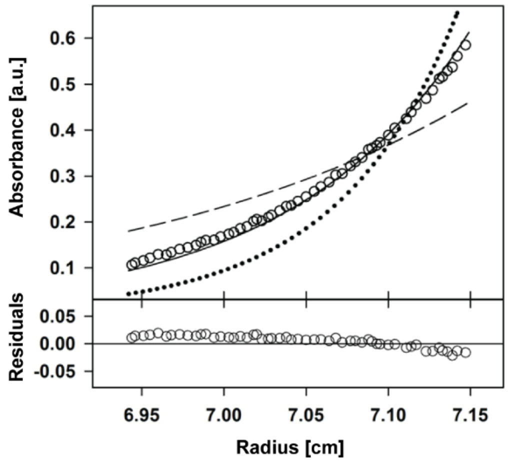


***__Supplementary Figure 1. Characterization of the ZapD dimer and ZapD-FtsZ-GDP interaction by Analytical ultracentrifugation (AUC).__*** **a** Sedimentation coefficient distribution, c(*s*), obtained by sedimentation velocity (SV) with (**I**) ZapD at different protein concentrations at working conditions (50 mM KCl, 50 mM Tris-Cl, 5 mM $MgCl_2$ pH 7). More than 95% of the sample sedimented as single species with an experimental sedimentation coefficient of 3.5 S once corrected to standard conditions (s20, w: 3.6 S ± 0.1). It matched with the expected value for ZapD dimers. (**II**) Sedimentation coefficient distribution, c(*s*), obtained by SV of FtsZ at 15 µM at working conditions (50 mM KCl, 50 mM Tris-Cl, 5 mM $MgCl_2$ pH 7). (**III**) Sedimentation coefficient distribution, c(*s*), obtained by SV of the interaction of 15 µM FtsZ and increasing concentrations of ZapD in physiological glutamate-acetate buffer. There is an axis break at 1.6 to highlight protein complexes. (**IV**) Represents the global multi-wavelength (280 and 250 nm)

analysis of the sedimentation coefficient distribution for FtsZ-ZapD complexes obtained by SV and decomposition into component sedimentation coefficient distributions $c_k(s)$ for FtsZ (solid trace) and ZapD (dashed trace) using FtsZ:ZapD initial ratio of 1:4. Numbers over the peaks correspond to the FtsZ:ZapD molar stoichiometry observed for each complex. **b** Concentration gradient obtained by sedimentation equilibrium (SE) of ZapD at 10 µM. Experimental data (empty circles) are shown together with the best-fit analysis corresponding to ZapD dimer (solid line), monomer (dashed line) and trimer (dotted line). The lower plot represents the residuals of the fitting of the experimental data to the dimeric model. ZapD was shown as a stable dimer, showing a buoyant mass of 14,415 Da, corresponding to a molecular mass of 56200 ± 400 Da, considering the partial specific volume calculated from the ZapD amino acid sequence.

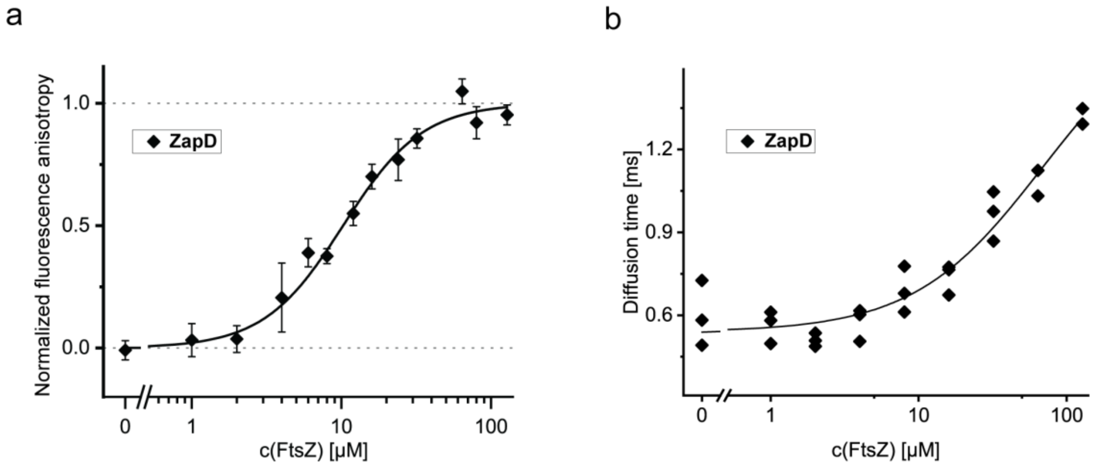


***__Supplementary Figure 2. Biochemical characterization of the ZapD-FtsZ-GDP interaction by FCS and fluorescence anisotropy.__*** **a** Interaction curve of FtsZ-GDP and ZapD by fluorescence anisotropy. The fluorescence anisotropy measurement of 5 µM ZapD supplemented with 150 nM ZapD-ATTO 647N at increasing concentrations of FtsZ demonstrated a direct interaction between both proteins with an apparent Kd of 10.26 ± 1.07 (SE). Samples were in working conditions (50 mM KCl, 50 mM Tris-Cl, 5 mM $MgCl_2$ pH 7) and the error bars represent the standard deviation among 3 independent samples. **b** Fluorescence correlation microscopy (FCS) analysis of increasing concentrations of FtsZ-GDP with 5 µM of ZapD supplemented with 100 nM ZapD-ATTO 647N in working conditions. The FCS analysis demonstrated an increasing diffusion time of ZapD along with the FtsZ concentration as result of higher proportion of ZapD bound to FtsZ. Negative controls were measured with either FtsZ or ZapD alone. The plotted line is only to guide the eye. Each condition was measured in three independent samples.

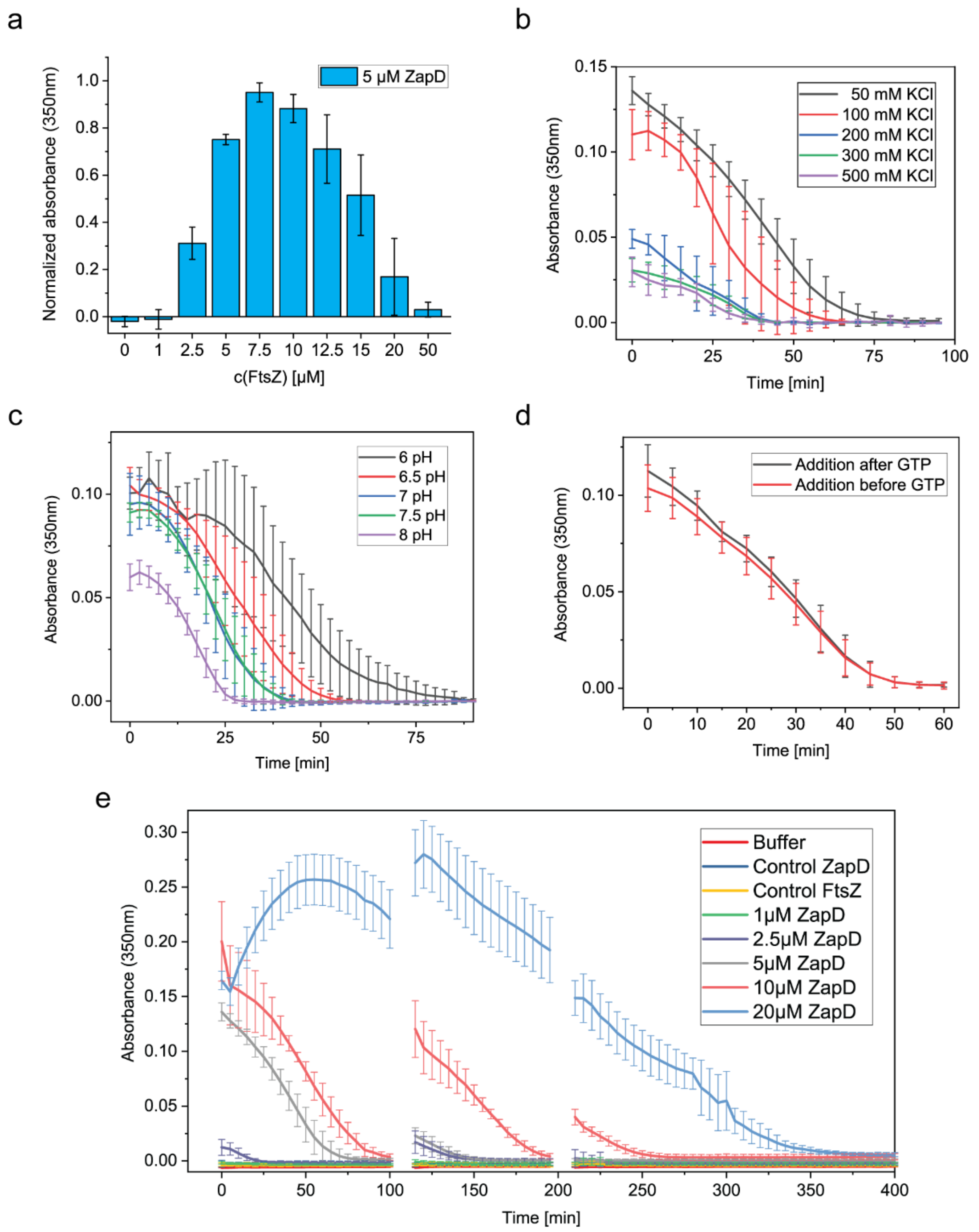


***<u>Supplementary Figure 3. Biochemical characterization of the ZapD and FtsZ bundles by Turbidity at 350 nm.</u>*** The characterization of the FtsZ bundles resulting from the interaction with ZapD was made by measuring the turbidity of the samples at 350nm in a plate reader. The mean value and standard deviation are the result of 3-6 independent samples. The signal from the blanks was collected before polymerization of FtsZ and subtracted for further measurements. Control of ZapD and FtsZ polymers independently did not show any significant difference with the blank after GTP addition. The working buffer was used unless it is specified in the legend (50 mM KCl, 50 mM Tris-Cl, 5 mM $MgCl_2$ and pH 7). The absorbance was measured every 5 min for 100 min. **a** FtsZ bundling at different concentration of FtsZ with 5 µM ZapD and 1 mM GTP in working conditions. In this case, the signal from FtsZ bundles was normalized to the maximum absorbance achieved (7.5 µM FtsZ) to facilitate the interpretation of the results. The signal plotted corresponds to 5 min after the addition of GTP. There was a clear decrease in the signal obtained at high concentration of FtsZ due to the low protein ratio ZapD/FtsZ. **b** Assembly of FtsZ bundles at different ionic strength conditions using KCl. The concentration of FtsZ and ZapD was 5 µM for both proteins, adding 1 mM GTP in the working buffer supplemented with different concentrations of KCl (50 - 500 mM KCl). Higher ionic strength in the buffer reduced the amount or size of bundles formed in solution, as the FtsZ-ZapD interaction is decreased. **c** Effect of pH in the FtsZ bundling process. The protein concentration used was 5 µM of ZapD and FtsZ with the addition of 1 mM GTP in the working buffer at different pH (6.5 - 8 pH). Lower pH enhanced the amount of FtsZ bundles promoted by ZapD, as lateral FtsZ-FtsZ interactions are promoted at lower pH. **d** Analysis of FtsZ bundling over time when ZapD was mixed with FtsZ prior or after the GTP addition and polymerization. The concentration of FtsZ and ZapD was 5 µM and 1 mM GTP in working buffer. ZapD mixed with FtsZ before or after GTP addition did not show significant difference in the formation of FtsZ bundles. **e** Measurement of turbidity over time of 5 µM FtsZ in the presence of ZapD at increasing concentrations. Both proteins were mixed prior to the addition of 1 mM GTP. A decay in the signal can be observed over time, highlighting the dynamic nature of the FtsZ bundles. After 100 min, extra 1 mM GTP was added and FtsZ bundles were reassembled, showing lower increase of the signal due to a higher concentration of GDP in the samples, likely sifting the reaction. The addition of 1 mM GTP was done twice after 100 min.

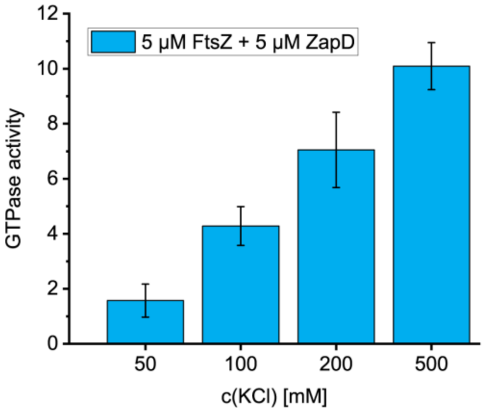


***Supplementary Figure 4. Ionic strength in the buffer lowers the effect of ZapD over FtsZ GTPase activity.*** GTPase activity of FtsZ in the presence of equimolar concentration of ZapD (5 µM) after addition of 1 mM GTP at different ionic strength conditions (50 – 500 mM KCl, 50 mM Tris-Cl, 5 mM $MgCl_2$ pH 7). The mean value and SD of the FtsZ GTPase activity correspond to 3 independent replicates. The GTPase activity was measured as result of the Pi released from GTP consumption. The units are Mol GTP consumed per Mol FtsZ per min. Moderate salt concentrations (100-200 mM) usually support optimal FtsZ assembly and GTP hydrolysis.

a

| ZapD [µM] | ZapD/FtsZ [mol/mol] |
|---|---|
| 0 | - |
| 1 | <0.05 |
| 5 | 0.3 ± 0.1 |
| 10 | 0.5 ± 0.1 |
| 30 | 1.1 ± 0.3 |

b

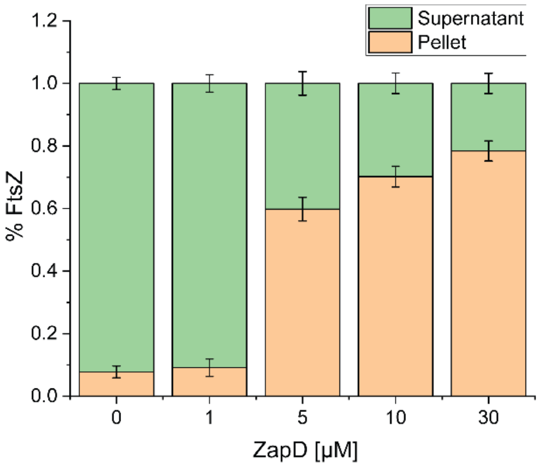


c

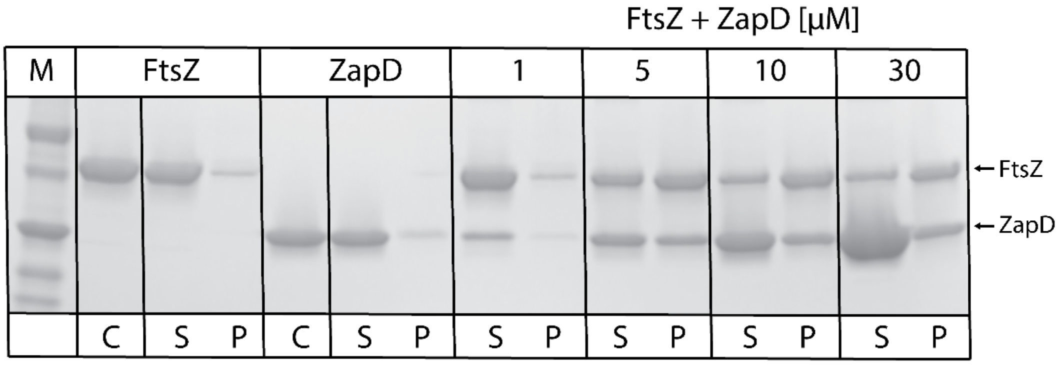


***<u>Supplementary Figure 5.</u> <u>ZapD binding to FtsZ Polymers via Sedimentation Assays.</u>*** **a** Analytical Sedimentation Velocity (SV) Assay. This table summarizes the binding of ZapD to FtsZ polymers, determined by the sedimentation velocity of FtsZ at 5 µM and increasing concentrations of ZapD (1 – 30 µM). The experiments were done at working conditions. **b** and **c** Pelleting Assays Using a Preparative Centrifuge. Samples of FtsZ (5 µM) and increasing concentrations of ZapD (1 – 30 µM) were sedimented at 10,000 rpm and the pellets and supernatants were subjected to SDS-PAGE (see panel c). The relative intensities in the electrophoresis allowed a qualitative estimation of the concentrations of FtsZ in the supernatant and pellet. **b** Shows the estimation of FtsZ in the pellet and supernatant. The protein concentrations and buffer conditions used in this assay were the same as those in the analytical SV assay. **c** *T*he samples loaded in the SDS gel from left to right include FtsZ at 5 µM, ZapD at 10 µM, and FtsZ in the presence of 1, 5, 10, or 30 µM ZapD. The molecular weight markers from top to bottom are 50, 37, 25, 20, and 15 kDa. "C" denotes the control sample without centrifugation, while "S" represents the supernatant, and "P" indicates the pellet.

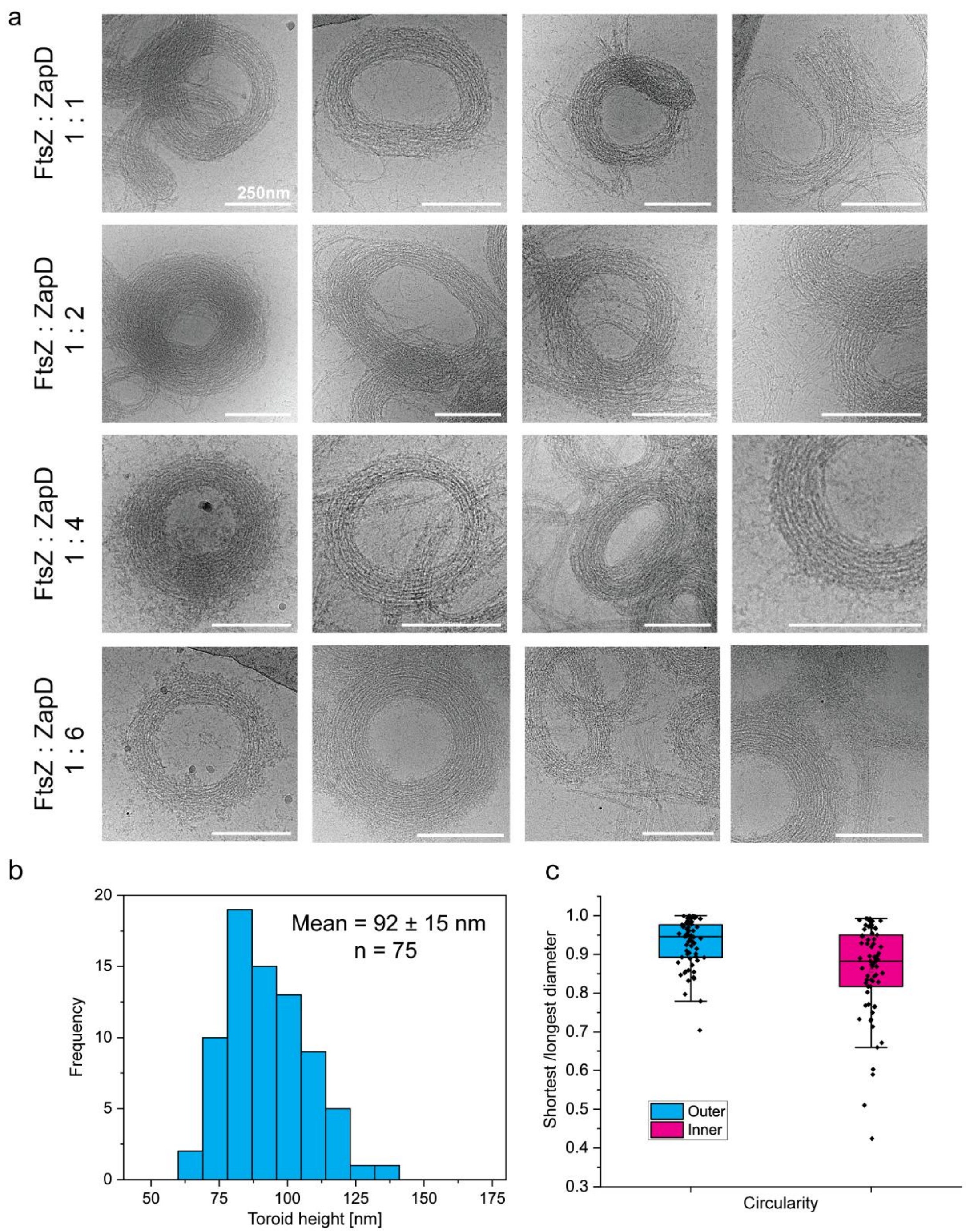


***Supplementary figure 6. FtsZ toroids and bundles formed at different protein ratios.*** **a** Cryo-EM images of FtsZ bundles and toroids promoted by interaction with ZapD at different protein ratios. From top to bottom, the protein ratios used in the samples were 1:1, 1:2, 1:4, 1:6 for FtsZ at 10 µM and ZapD concentrations ranging from 10 to 60 µM. These ratios do not refer to the stoichiometry of the proteins forming the structure. Samples were plunge frozen two minutes after the addition of 1 mM GTP. In these samples, similar toroidal structures and bundles were observed regardless of the protein ratio, including spirals and curved bundles. Scale bars are 250 nm. **b** Distribution of toroids heights measured from cryo-ET data. Cross-section of the toroids in the XZ plane were used to measure the height of each toroid. Distance measurements were taken from each toroid in the middle of the bundle at the four cardinal points to assure the reliability of the results. The mean value and SD are shown in the graph. **c** Circularity of FtsZ toroids. For each toroid, the shortest and largest distances of the outer and inner diameter were measured from the cryo-EM images. The division between the shortest and longest diameter provided the circularity of the FtsZ toroids. The mean value and SD are shown in the graph. The circularity of the inner and outer diameter is significantly different (T-test t(4.2), p-value <0.001).

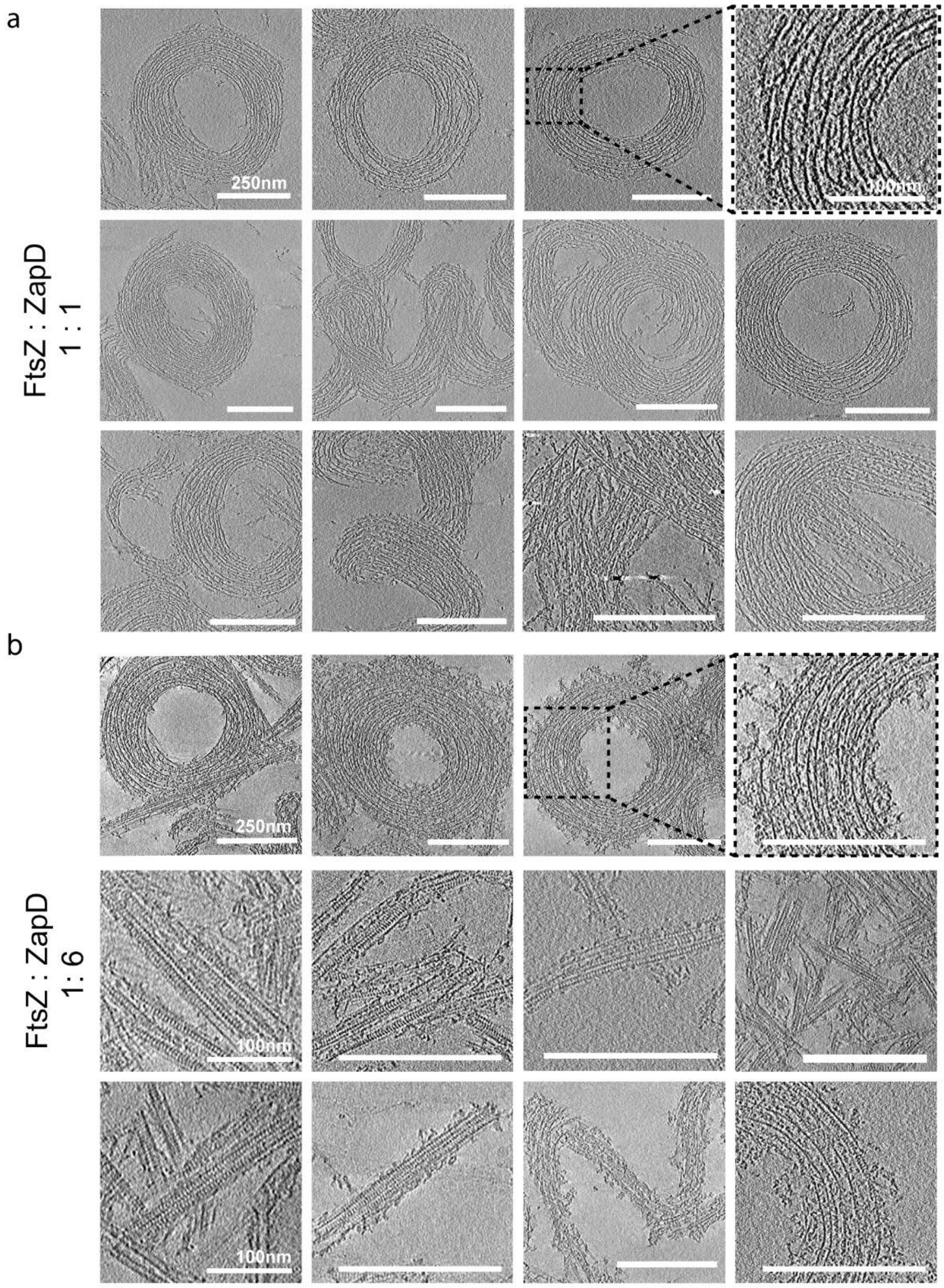


***Supplementary figure 7. FtsZ structures found at low and high ZapD concentrations.*** Representative tomographic slices of FtsZ structures and bundles promoted by ZapD at different protein ratios under working conditions. The images are an average of five 0.86 nm slices from the reconstructed tomographic volume (total thickness of 4.31 nm) to enhance the signal to noise ratio of the images. Scale bars are 250 nm unless they are labelled with 100 nm. Samples were plunge frozen 2 min after triggering polymerization. **a** Equimolar concentrations of FtsZ and ZapD (10 µm) in the sample formed toroids and bundles after 1 mM GTP addition. **b** High concentration of ZapD (60 µm) promoted the formation of straight FtsZ bundles, although they coexisted with scarce amount of FtsZ toroids and double filaments showing striated patterns.

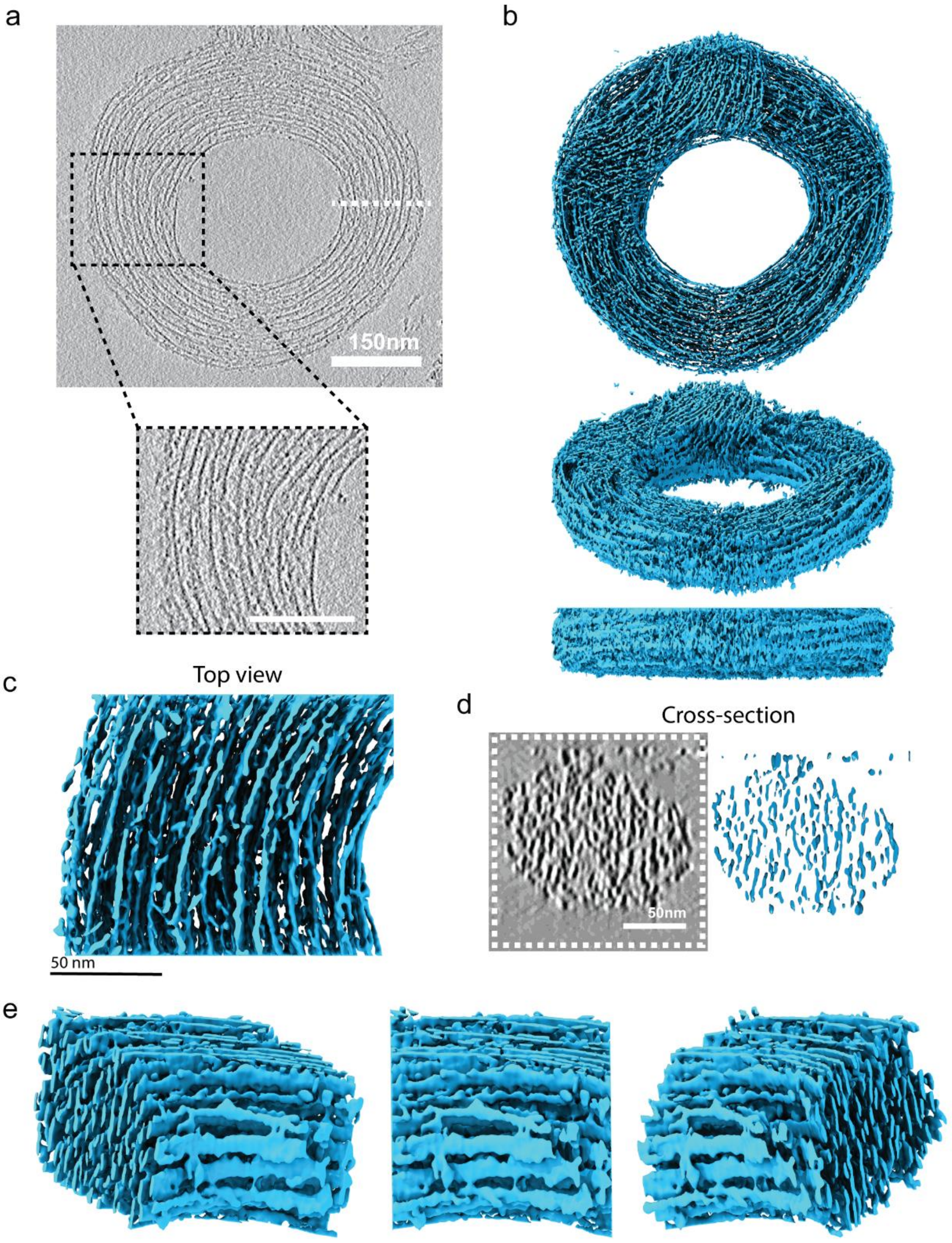


***<u>Supplementary Figure 8. Segmentation of a FtsZ toroid imaged using cryo-ET</u>.*** **a** Tomographic slice of a toroidal FtsZ structure. The image is an average of five 0.86 nm slices from the reconstructed tomographic volume (total thickness of 4.31 nm). FtsZ and ZapD were added to the sample at equimolar concentrations (10 µm) before the addition of 1 mM GTP. The white dotted line represents the localization of the cross-section shown in **e**, while the black dotted square shows the area of a close-up view shown below the image. The scale bar of the zoomed area is 50 nm. **b** Corresponding segmentation of the toroid shown in **a.** The isosurface of the toroid was extracted from the denoised tomographic volume and positioned in different views: front (top), side (middle) and lateral (bottom). The curved filaments observed in the upper part of the toroid correspond to a bundle that was located in the upper layer of the toroid. It has been erased to make the toroid easier to see. The toroid has a diameter of ~581 nm and a height of ~119 nm. **c** Segmentation of the zoomed area shown in **a**. Top view of the isosurface from the toroidal structure. **d** Cross-section of the toroid in the XZ plane (left; plane indicated by the white dotted line in **a**) and corresponding isosurface (right). The tomographic slice is the average of nine tomographic slices. **e** Different views of the isosurface of the toroidal structure. The network of filaments shown corresponds to the image shown in **c** in different views. The segmentation shown has a width of 116 nm x 150 nm and a height of 82 nm.

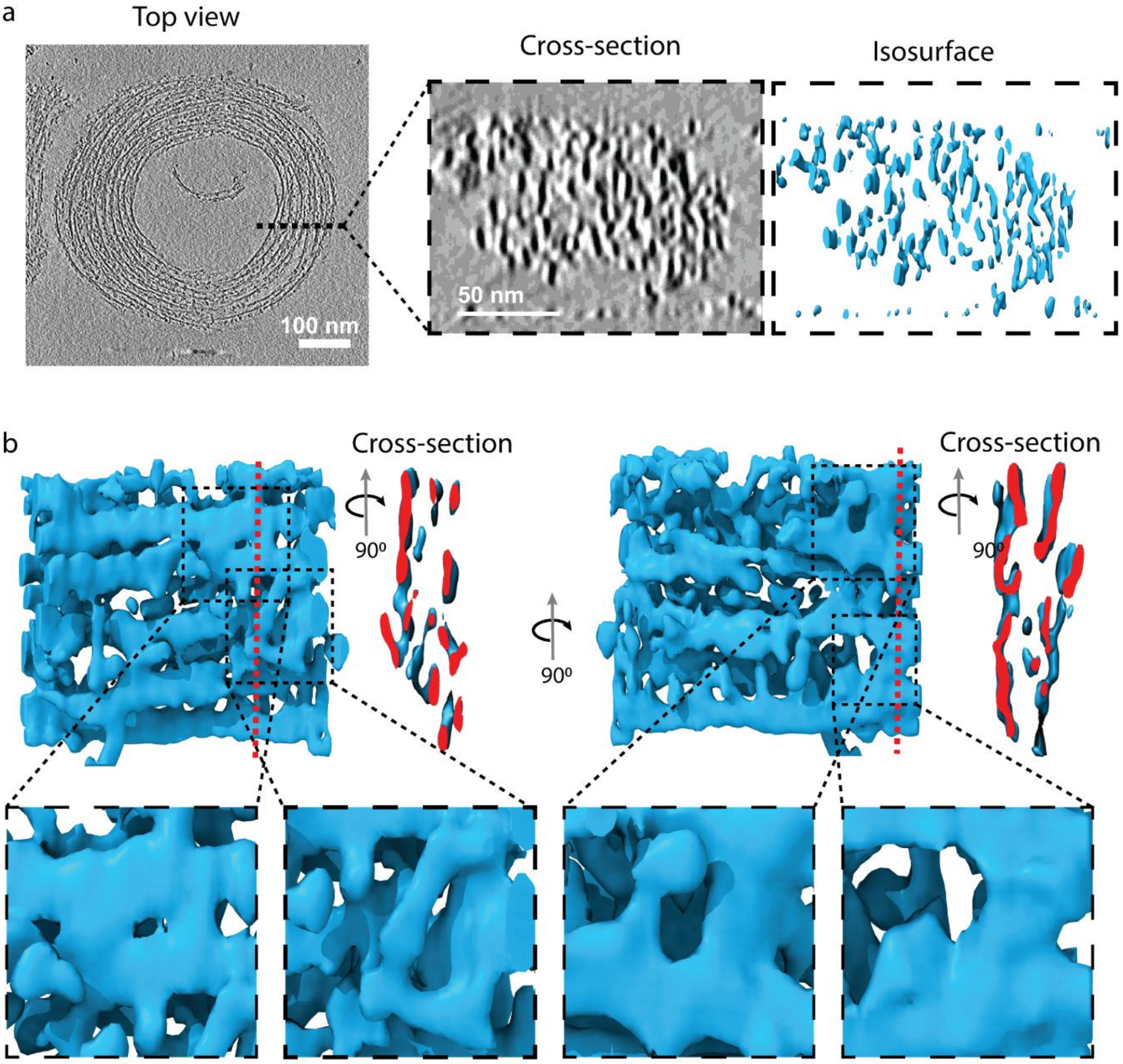


***__Supplementary Figure 9. Cross-section of the toroid showing the elongated structures in the Z-axis__.*** **a** Tomographic slice of a toroid. The image is an average of five 0.86 nm slices from the reconstructed tomographic volume (total thickness of 4.31 nm). The tomographic slice in the XZ plane shows a cross-section of the toroid (middle). This image is the average of nine tomographic slices (total thickness of 7.74 nm) from the denoised tomogram. Corresponding isosurface of the same area (right). **b** Different views of the isosurface of a filament network extracted from a toroid. The red dotted lines indicate the area that was selected for the cross-sections. Black dotted squares show closer views of the connections between filaments in Z-plane. The segmentation shown has a width of 28 nm x 37 nm and a height of 56 nm.

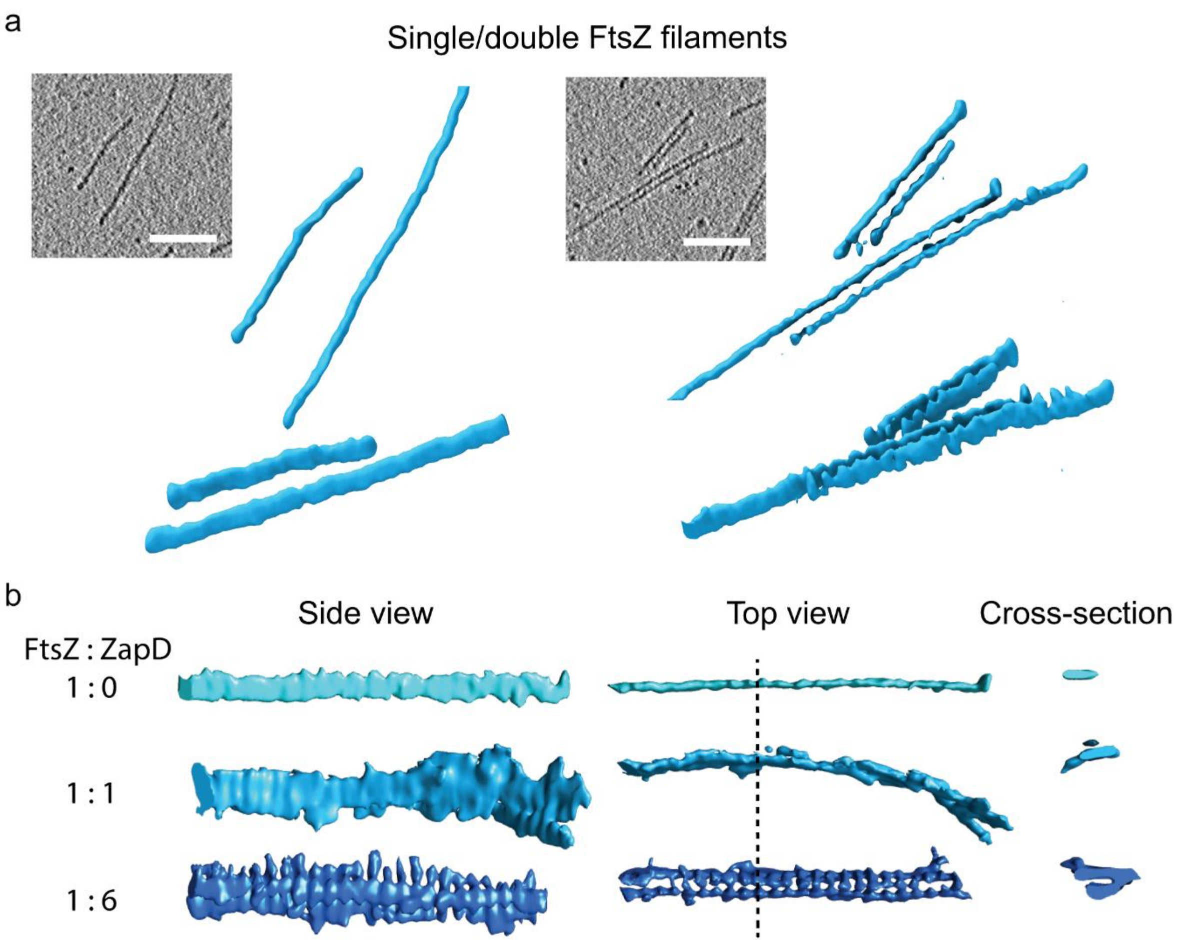


***__Supplementary figure 10. Segmentation of FtsZ filaments__.*** **a** Isosurface of single and double FtsZ filaments in the absence of ZapD extracted from the denoised tomographic volume. FtsZ filaments observed from cryo-ET samples in the presence of 2 mM GTP. The single and double filaments were extracted from the tomographic slice shown to the left side of each example. Top and side views of the segmented filaments are shown. Scale bars represent 50 nm. **b** Comparison of isolated FtsZ filaments from the following samples: (top) absence of ZapD, (middle) a toroidal structure at equimolar concentrations of FtsZ and ZapD (10 µM) and (bottom) a straight bundle at high concentration of ZapD (60 µM). The protein ratios added to the sample in each case are shown in the left side of each example. Side and top views of the three segmented filaments are shown to the left and middle columns, while a cross-section of the filaments is shown in the right column. The cross-section corresponds to the dotted line shown at the side top view. Different shades of blue distinguish the three conditions analyzed. The presence of extra densities decorating the FtsZ filaments strongly suggest that these are ZapD proteins decorating and connecting the filaments. These densities generate an elongation in Z beyond the missing wedge, which is clearly visible.

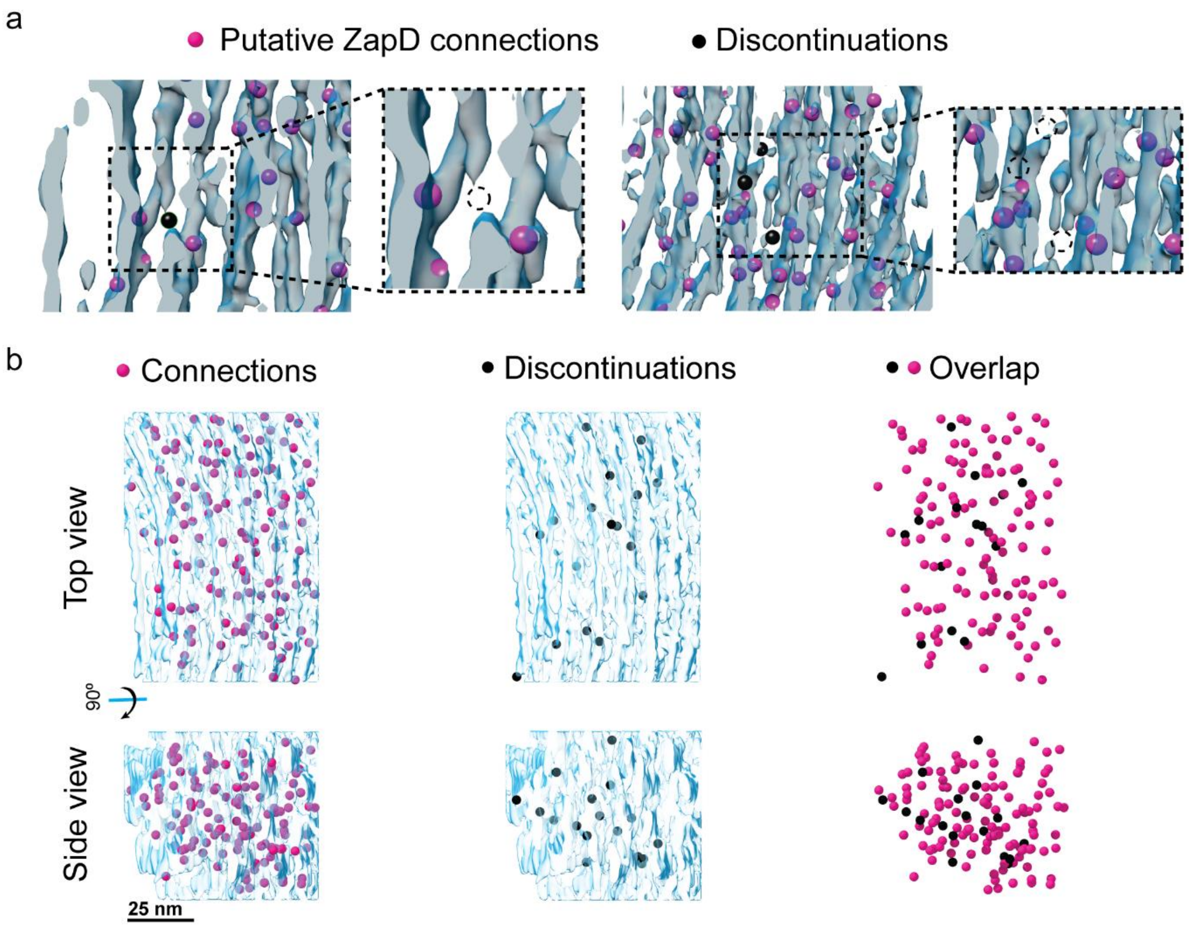


***Supplementary figure 11. The FtsZ toroid is formed by short and discontinuous filaments crosslinked by ZapD.*** **a** Image of the isosurface of the network of filaments shown in Fig. 4a. A black marker (sphere) is positioned at each discontinuity or termination of FtsZ filaments (blue) within the toroid. Magenta markers were placed at each clear connection between FtsZ filaments to analyze their presence in areas close to discontinuities. Closer inspection of the image, framed by a dotted black line, shows the termination and interruption of some FtsZ filaments. The black marker has been replaced by a dotted circle to show the absence of filaments in this area. The images on the left and right are just different areas of the same network of filaments within the toroid. **b** Images on the left and in the middle represent the spatial 3D localization of discontinuities (black) and connections (magenta) of FtsZ filaments in the analyzed area of the toroid shown in (Fig. 4a) (blue filaments). A top and side view of the structure show the localization of the discontinuities and connections. The overlap of the two (right) shows no significant colocalization or correlation between them. The segmentation shown has a width of 73 nm x 101 nm and a height of 64 nm.

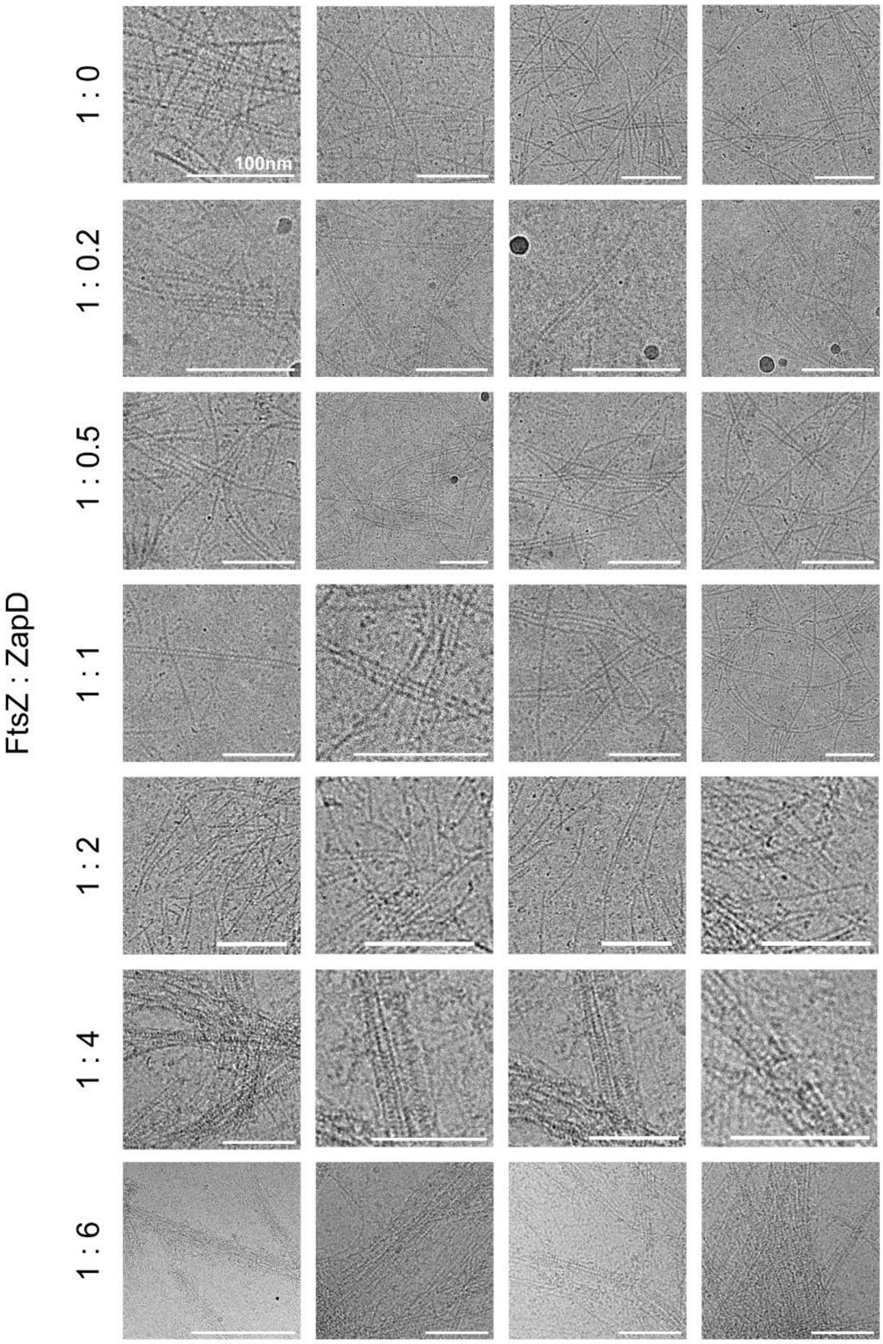


***Supplementary figure 12. FtsZ single and double filaments found at different FtsZ and ZapD ratios.*** Cryo-EM images of single and double filaments found together with toroids and bundles in samples containing FtsZ and ZapD at different protein ratios in the sample. The FtsZ concentration was 10 µM, while ZapD was present at increasing concentrations (from top to bottom 0, 2.5, 5, 10, 20, 40 and 60 µM), mixed before the addition of 1 mM GTP. These ratios refer to the initial protein concentration added and not to the stoichiometry of the two proteins forming the structure. In the cryo-EM samples, only single and double filaments were found, as shown in samples from 1:0 to 1:1 protein ratio. In these filaments, we could not discard or confirm the presence of ZapD connecting the FtsZ filaments. They were not easily distinguished from double filaments formed in the absence of ZapD as a consequence of weak FtsZ-FtsZ lateral interactions. In the case of saturation of ZapD in samples containing 1:4 or 1:6, double FtsZ filaments and straight bundles with a characteristic striped pattern perpendicular to the FtsZ filaments were found. Scale bars are 100 nm.

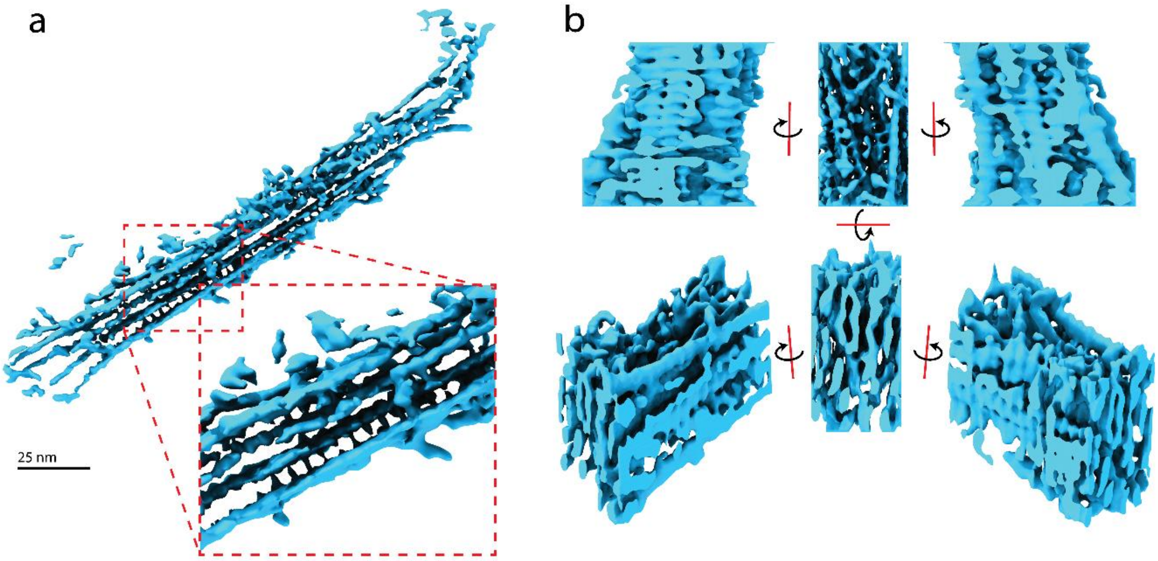


***Supplementary Figure 13. Segmentation of straight bundles imaged by cryo-ET.*** Segmentation of straight bundles resulting from the interaction of FtsZ with high concentrations of ZapD under working conditions (50 mM KCl, 50 mM Tris-Cl, 5 mM $MgCl_2$ pH 7). The concentrations of FtsZ and ZapD were 10 µM and 60 µM, respectively. 1 mM of GTP was added to induce polymerization and bundle formation. **a** and **b** show the isosurface of two straight bundles from the denoised tomographic volume. The entire volume has been colored in blue without differentiating ZapD connections to facilitate the interpretation. **b** shows different views of a cropped area within a straight bundle. The segmentation shown in **a** has a width of 270 nm x 175 nm and a height of 72 nm. Segmentation in b has a width of 36 nm x 65 nm and 60 nm in height.

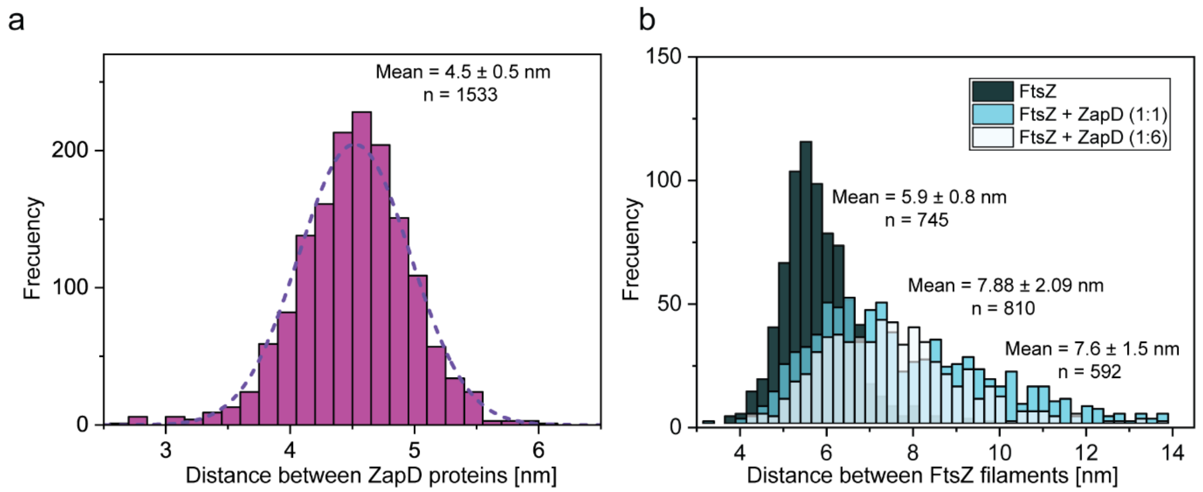


***<u>Supplementary figure 14. Distance between FtsZ filaments and ZapD-associated filaments.</u>*** **a** Distribution of the distance between two ZapD proteins in a straight bundle under ZapD saturation (1:6 protein ratio). The distance between two ZapD proteins was measured from tomographic images obtained in samples containing FtsZ (10 µM) in the presence of ZapD at high concentration (60 µM) in the presence of GTP. The distribution obtained fitted a normal distribution. The mean value and SD are shown in the graph. The regular distance found between the ZapD proteins connecting the filaments could indicate the presence of one ZapD dimer per FtsZ monomer forming the filament. **b** Distribution of the distance between two FtsZ filaments (5.9 ± 0.8 nm) in the absence or presence of ZapD forming toroidal structures (1:1) (7.88 ± 2.09 nm) or straight bundles (1:6) (7.6 ± 1.5 nm). The distance between filaments was measured from the cryo-EM and cryo-ET images and plotted in the graph as a distribution of distances. The three distributions were obtained from images of >3 independent samples. The mean value and SD are shown in the graph for each condition. The presence of ZapD crosslinking filaments inreases the distance between them compared to weak FtsZ-FtsZ interactions. However, no significant differences were found between equimolar and saturation levels of ZapD.

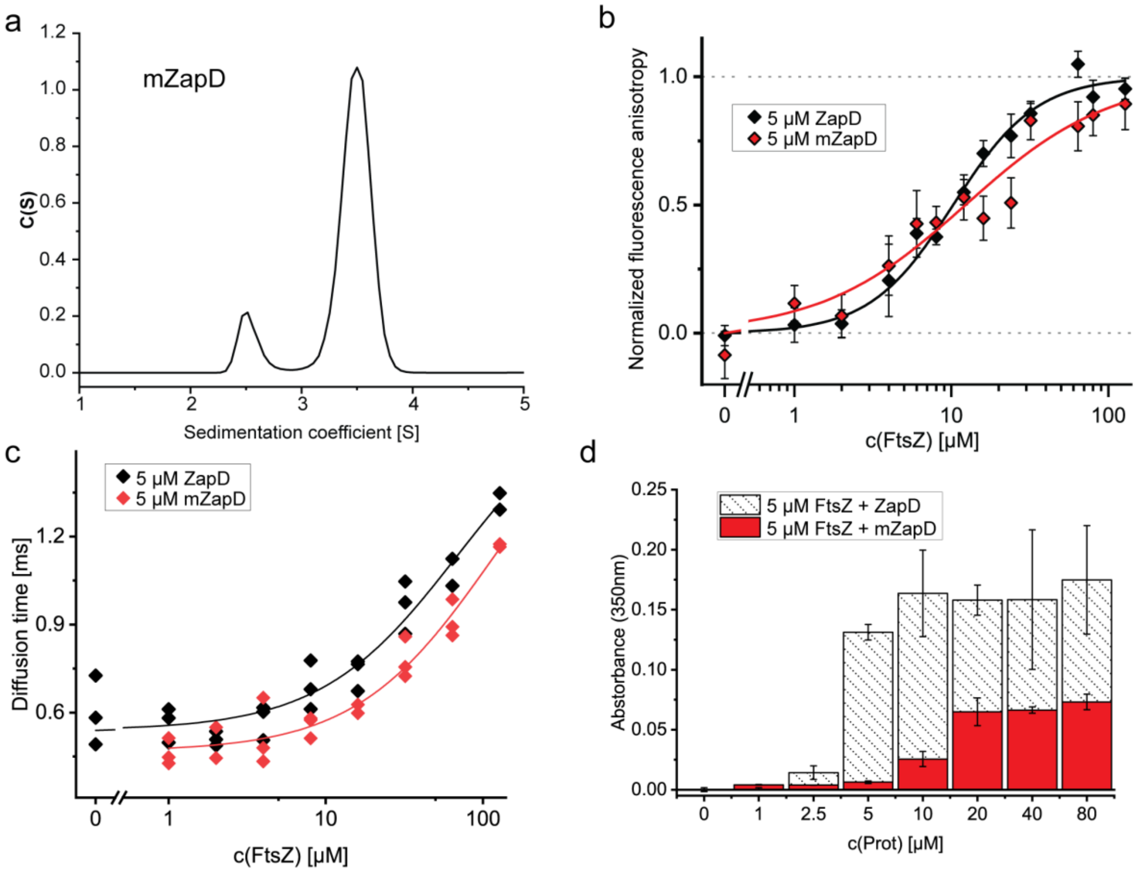


***<u>Supplementary figure 15. mZapD binds FtsZ-GDP and can promote bundling of FtsZ filaments.</u>*** **a** Sedimentation coefficient distribution, c(*s*), obtained by sedimentation velocity (SV) with mZapD at different concentrations under working conditions (50 mM KCl, 50 mM Tris-Cl, 5 mM $MgCl_2$ pH 7). mZapD sedimented mostly as a single species with an experimental sedimentation coefficient of 3.5 S when corrected to standard conditions (s20, w: 3.6 S ± 0.1). However, species at around 2.5 S appear at ~10%, in contrast to ZapD, demonstrating the presence of monomers and confirming the weakening of the dimerization interface. **b** Molecular interaction between FtsZ-GDP and ZapD or ZapD mutant (mZapD) measured by fluorescence anisotropy and compared with the results shown in Supplementary Fig. 2a for ZapD and FtsZ-GDP. 5 µM of mZapD was supplemented with 150 nM of mZapD-ATTO 647N and FtsZ in increasing concentrations. mZapD (red) also binds FtsZ-GDP with an apparent Kd of 12.42 ± 5.45 (SE). The x-axis is expressed in log units. The mean value and standard deviation are the result of 3 independent samples. **c** Fluorescence correlation spectroscopy (FCS) analysis of increasing concentrations of FtsZ-GDP with 5 µM of ZapD or mZapD supplemented with 100 nM ZapD or mZapD chemically labelled with ATTO 647N. The plot corresponding to ZapD was the same as shown in Supplementary Fig. 2b and it is included here to facilitate the interpretation of the results. The x-axis is expressed in log units. The diffusion curve showed an increasing diffusion time of mZapD with increasing FtsZ concentration, supporting a direct interaction of this protein with FtsZ-GDP. The plotted line is to guide the eye only. Each condition was measured in three independent samples. **d** Turbidity signal of FtsZ and ZapD by measuring absorbance at 350 nm after 5 min of GTP addition. The concentration used was 5 µM FtsZ with increasing concentrations of ZapD (0 - 80 µM) and 1 mM GTP in working buffer. Data represents the mean vand standard deviation of >3 independent samples. The signal shown for ZapD is the same as that shown in Fig. 1b and it is included here to aid interpretation.

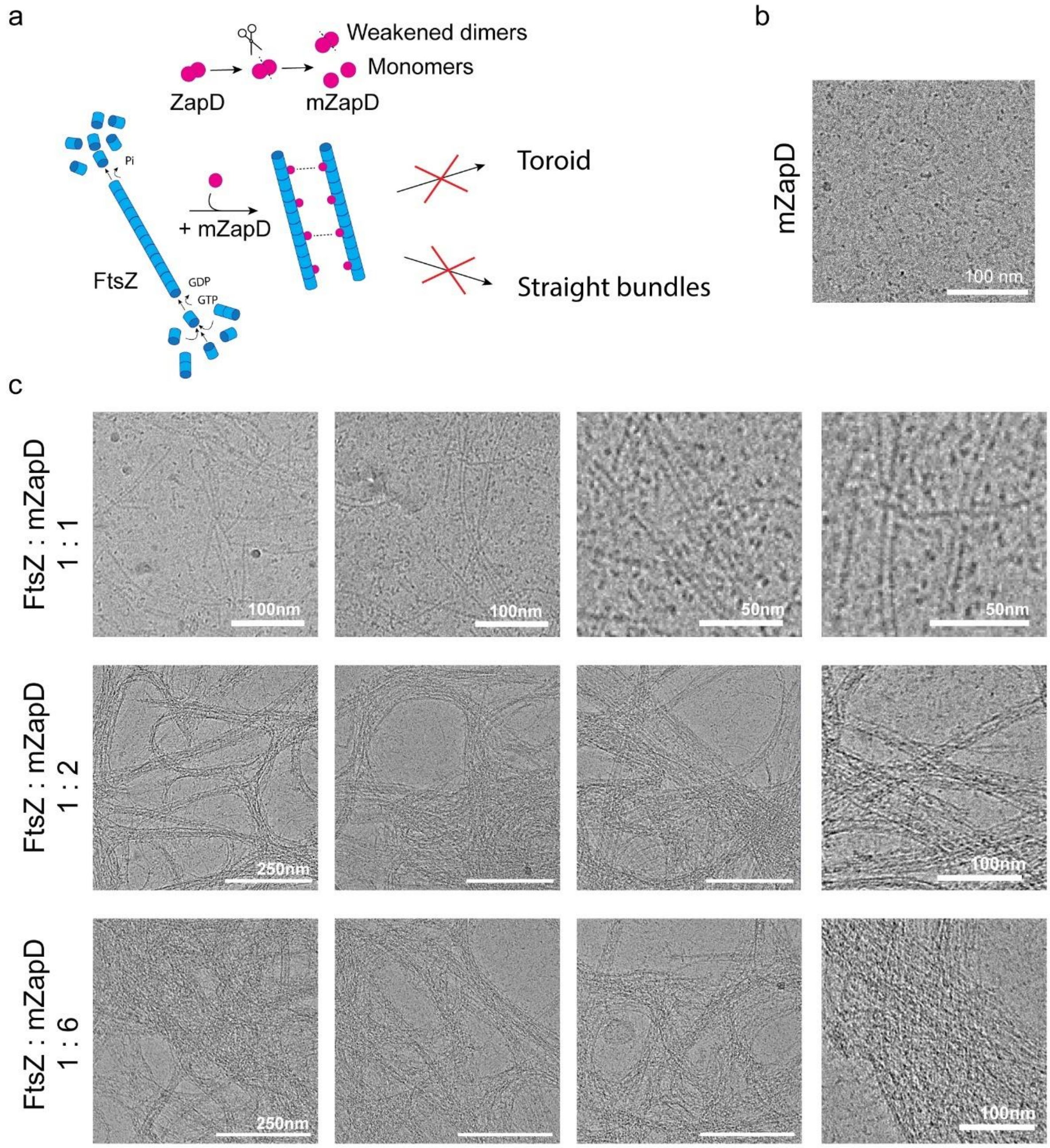


***<u>Supplementary figure 16. mZapD can bundle FtsZ without promoting toroids.</u>*** **a** Proposed schematic of the interaction mechanism between FtsZ-GTP and the mZapD protein. The mZapD protein is mutated at the dimerization site to weaken dimerization and favor monomerization of the protein. Thus, the interaction of dynamic FtsZ filaments with mZapD monomers is weakened and the stabilization of toroidal structures or straight bundles is not possible, while dynamic bundles can still be formed. **b** Cryo-EM image of 10 µM mZapD. mZapD is homogeneously distributed and it did not aggregate under our working conditions. The scale bar is 100 nm. **c** Representative cryo-EM images of FtsZ and mZapD at increasing concentrations of mZapD and 1 mM GTP under working conditions. The ratios shown on the left side of the images refer to the initial added protein concentration. At equimolar concentrations of FtsZ and ZapD (10 µM), no FtsZ bundle were found and only single and double FtsZ filaments were observed (top). At higher concentrations of mZapD (20 and 60 µM, middle and bottom rows, respectively), FtsZ bundles were formed, although no toroids or straight bundles were observed in these samples. The absence of these structures suggested that a stable dimer is required to form them. Scale bars are 250 nm unless 100 nm or 50 nm labels are shown.

***<u>Supplementary Table 1. List of primers.</u>***

| Name | Sequence (5’ to 3’) |
|---|---|
| ZapD-R20A | GAAAAAATGCGTACATGGCTG**GCT**ATTGAGTTTT |
| ZapD-R116A | GGCAATTTCTGCGTGAAGAT**GCT**TTGATTGCTC |
| ZapD-H240A | CTTTGATTTACCTACATTG**GCT**ATTTGGCTGC |
